# Identification of differential gene expression pattern in lens epithelial cells derived from cataractous and non-cataractous lenses of Shumiya cataract rat

**DOI:** 10.1101/2020.05.19.103879

**Authors:** Hidetoshi Ishida, Teppei Shibata, Yuka Nakamura, Yasuhito Ishigaki, Dhirendra P Singh, Hiroshi Sasaki, Eri Kubo

**Affiliations:** Department of Ophthalmology, Kanazawa Medical University, Kanazawa, Ishikawa, Japan; Medical Research Institute, Kanazawa Medical University, Kanazawa, Ishikawa, Japan; Department of Ophthalmology, University of Nebraska Medical Center, Omaha, Nebraska, United States of America

**Keywords:** Shumiya cataract rat, microarray, *Dnase2b*, *HspB1*, *MIP*, γ-crystallin

## Abstract

The Shumiya cataract rat (SCR) is a model for hereditary cataract. Two-third of these rats develop lens opacity within 10-11-weeks. Onset of cataract is attributed to the synergetic effect of lanosterol synthase (*Lss*) and farnesyl-diphosphate farnesyltransferase 1 (*Fdft1*) mutant alleles that lead to cholesterol deficiency in the lenses, which in turn adversely affects lens biology including the growth and differentiation of lens epithelial cells (LECs). Nevertheless, the molecular events and changes in gene expression associated with the onset of lens opacity in SCR is poorly understood. In the present study, a microarray-based approach was employed to analyze comparative gene expression changes in LECs isolated from the pre-cataractous and cataractous stages of lenses of 5-, 8- and 10-week-old SCRs. The changes in gene expression observed in microarray results in the LECs were further validated using real-time PCR and western blot analyses. Lens opacity was not observed in 5-week-old rats. However, histological analysis revealed mild dysplasia in the anterior suture and poorly differentiated fiber cells at the bow region. Expression of approximately 100 genes, including major intrinsic protein of lens fiber (*MIP* and Aquaporin0), deoxyribonuclease II beta (*Dnase2b*), heat shock protein B1 (*Hspb1*), and crystallin γ (*γCry*) B, C, and F were found to be significantly downregulated (0.07-0.5 fold) in rat LECs derived from cataract lenses compared to that in noncataractous lenses (control). Thus, our study aimed to identify the gene expression patterns during cataract formation in SCRs, which may be responsible for cataractogenesis in SCR. We proposed that mutation in lanosterol synthase was responsible for the downregulation of genes associated with lens fiber differentiation, which in turn leads to the formation of cataract in these rats. Our findings may have wider implications in understanding the effect of cholesterol deficiency and the role of cholesterol-lowering therapeutics on cataractogenesis.

## Introduction

Age related eye disease is a serious public health issue and age-related cataract is the leading cause of blindness worldwide [1]. Currently, surgery is the only treatment for cataract. It has been reported that if the progression of cataract is delayed by 10 years, the huge expense associated with surgical intervention can be reduced [2]. Cataracts are caused by the degeneration of lens protein called crystallin. Lenses with almost no protein turnover are susceptible to ultraviolet rays, oxidative stress, and glycative stress. These stressors damage protein integrity and function leading to denaturation and aggregation of lens protein, which in turn results in lens opacification [3, 4]. Recent studies have shown that oxidative stress controls various cellular processes associated with cell survival, such as cell proliferation, differentiation, aging, and cell death. It promotes cell apoptosis and senescence and is also associated with many diseases [5, 6]. Oxidative stress and reactive oxygen species (ROS) are a major cause of age-related eye diseases, and diets rich in fruits, vegetables, vitamin C, zeaxanthin, lutein, and multivitamin-mineral supplements are recommended for cataracts and age-related macular degeneration (AMD) [7-10]. Moreover, several risk factors such as advancing age, genetic predisposition, oxidative stress and external and internal factors causing adversely on lens homeostasis have been implicated to contribute to the etiology of cataract formation.

Cataract is suggested to be a multifactorial disease associated with multiple etiological factors. To understand the molecular mechanism underlying cataract formation, animal models are a prerequisite. Shumiya cataract rats (SCRs) have been used as a model to study congenital cataract, exploring the initiation and development of the disease [7-9]. Lens opacity is observed in 66.7% of SCRs [10]. Onset of mature cataract in SCR occurs at around 10-11 weeks of age [10]. The vertebrate lens has a single layer of epithelial cells on its anterior surface. These cells are metabolic engines of lens and are responsible for maintaining its homeostasis and transparency. Any damage to lens epithelial cells (LECs) leads to cataractogenesis. Malfunctioning and aberrant differentiation of LECs contribute to cataractogenesis in SCRs [11] Another interesting aspect is that the onset of cataract occurs in lens epithelium in various types of cataract, such as selenite-induced cataract, UV exposed cataract (*in vitro*), diabetic cataracts in rats, and non-congenital cataracts in humans [12-16] The SCR is a model for hereditary cataract, but it also displays the features of oxidative stress-induced cataracts as its onset could be delayed by the administration of antioxidants [9] Moreover, the genetic basis of cataractogenesis in SCRs is associated with the combination of lanosterol synthase (*Lss*) and farnesyl diphosphate farnesyl transferase 1 (*Fdft1*) mutant alleles in the cholesterol biosynthesis pathway, lowering cholesterol levels in SCR lenses [16]. A previous study showed that inhibition of cholesterol synthesis inhibits cell proliferation in lens epithelial cells (LECs) [17], implicating that adequate cholesterol synthesis is required for maintaining the integrity and function of LECs, which in turn maintains normal lens biology and homeostasis. It has been found that mutations in *L*ss is one of the causes for the onset of human congenital cataracts [18, 19]. Furthermore, lens treated with lanosterol showed decreased opacity and increased transparency in case of canine cataract and reversed the aggregation of crystallin *in vitro* [18]. Hence, SCR is an appropriate model to study the mechanism underlying cataract formation and could help in the development of drugs for the treatment of cataract.

In this study, we investigated the gene expression changes in LECs of SCRs to analyze the mechanism underlying cataract formation and its association with *Lss* mutation and cholesterol deficiency. A comprehensive analysis of gene expression changes in LECs from SCRs was screened using DNA microarray analysis to identify genes associated with cholesterol deficiency. Our study may identify the various genes contributing to the onset of congenital cataract due to cholesterol deficiency. The outcomes of this study could contribute to the development of various therapeutic strategies to treat cataract.

## Materials and Methods

### Animals

All animal experiments were approved by the Committee of Animal Research at Kanazawa Medical University (Permission no. 2017-107) and were conducted in accordance with the Guide for the Care and Use of Laboratory Animals implemented by National Institutes of Health, the recommendations of the ARVO Statement for the Use of Animals in Ophthalmic and Vision Research, and the Institutional Guidelines for Laboratory Animals of Kanazawa Medical University. SCRs (SCR/Sscr: NBRP Rat No: 0823) were obtained from the National BioResource Project - Rat, Kyoto University (Kyoto, Japan).

We used 5-, 8- and 9-week-old SCRs in this study. All rats were provided ad libitum access to regular or experimental chow (Sankyo Labo Service, Tokyo, Japan). The animals were sacrificed by administration of a lethal dose of CO_2_. Cataractous (Cat+) and non-cataractous (Cat-) SCRs were distinguished via PCR using genomic DNA isolated from the tails of 4-week-old rats. The amplified products were then separated on a 15% gel via polyacrylamide gel electrophoresis (PAGE) to detect *Lss* mutation. The sequences of primers used to detect *Lss* mutation were as follows: 5’-GCACACTGGACTGTGGCTGG-3’ and 5’-GCCACAGCATTGTAGAGTCGCT-3’.

### RNA extraction

Total RNA from each LEC sample obtained from SCR was extracted using the miRNeasy Micro Kit (Qiagen, Valencia, CA) following the manufacturer’s protocol. Since RNA is extremely susceptible to degradation due to the ubiquitous presence of RNAses in the environment, purity and integrity of RNA were examined and validated as previously described [20]. Quality of total RNA was analyzed by evaluating the RNA integrity number (RIN) using Bioanalyzer RNA analysis (Agilent Technologies Japan Ltd., Tokyo, Japan). All RNA samples showed RIN > 9.0.

### Microarray analysis

Total RNA from each LEC sample obtained from 5-week-old cataractous (Cat+) and non-cataractous (Cat-) SCRs were used for the microarray analysis (n=1, each). All samples were processed for the microarray analysis as follows: For RNA labeling and hybridization, GeneChip(tm) WT Pico Reagent Kit (Applied Biosystems, ThermoFisher Scientific Tokyo, Japan) and GeneChip^®^ Rat gene 2.0 ST array (Affymetrix, ThermoFisher Scientific) were used according to the manufacturer’s protocol. Washes, scanning of the arrays, and analysis of scanned images were performed according to manufacturer’s instructions. Each chip was normalized by dividing the measurement of each gene by the measurement of the specific control or by average intensity in the single array. Normalized data were exported for subsequent analysis. Genes with a normalized ratio >2.0-fold or <0.5-fold were selected as significant genes using GeneSpring software package version 14.9 (Agilent).

### Real time quantitative reverse transcription PCR (RT-qPCR)

To measure the expression of rat deoxyribonuclease II beta (*Dnase2B*), heat shock protein B1 (*HspB1*), major intrinsic protein of lens (*MIP*) and crystallin, gamma D (*CryγD*) mRNAs, we conducted a relative quantification of mRNA using Prism7300 (Applied Biosystems, ThermoFisher Scientific). The comparative Ct method was used for relative quantification of mRNA expression. The PCR amplification was performed with TaqMan Universal PCR Master Mix. The probe mix containing the primers for *Dnase2B, HspB1, γCry*, and *MIP* was obtained from Thermo Fisher Scientific. All reactions were performed in triplicates. Differential expression for each gene was calculated using the comparative CT method using a pre-developed TaqMan Ribosomal RNA Control Reagent VIC probe as an endogenous control (ThermoFisher Scientific).

### Statistical analysis

The statistical analysis was performed for all experiments using Student’s t-test and/or one-way analysis of variance (ANOVA), as applicable. The data were presented as the mean ± standard deviation (SD). A significant difference between the control and treatment group was defined as a *p*-value <0.05 for two or more independent experiments.

## Results

### Lens opacity was observed in 5- and 10-week-old SCRs

In Cat-(noncataractous) rats, lens was clear, even at 10 weeks of age, but in case of Cat + (cataractous) rats, there was a mild posterior and cortical opacity was observed at 5 weeks of age, and severe cortical and nuclear opacity was observed at age 10 weeks (Fig. 1).

**Fig. 1:**
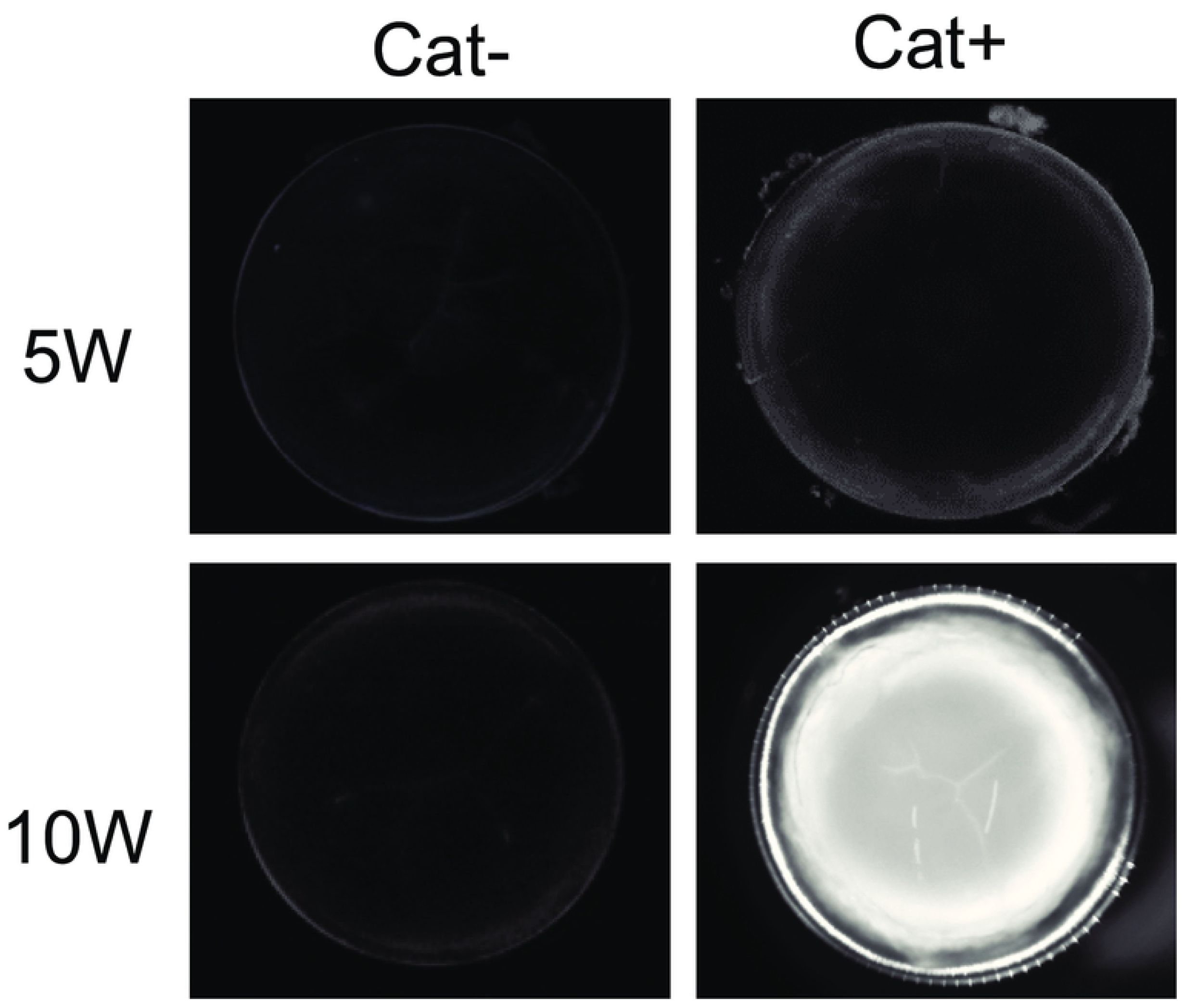
Observation of lens in 5- and 10-week-old Cat- and Cat+ SCRs. Lenses were extracted from 5 and 10-week-old SCRs and placed on glass bottom dishes containing Medium199 (Thermo Fisher Scientific). Photographs of the lens were acquired and recorded with a stereo microscope.

### Analysis of gene expression profile

As described in the Materials and Methods, LECs from, 5-week-old Cat+ and Cat-SCR were used for microarray analysis (n=1 in each group) s to screen genes associated with cholesterol deficiency and cataract. First, with 28,407 genes on the array, 110 genes were detected that showed a fold change <0.5 in the Cat+ group compared to those in Cat-, as shown in Fig. 2. In the Scatter plot, many genes were distributed in the lower part, and specifically, genes whose expression was downregulated could be detected using the scatter plot (Fig. 2). Fold changes (<0.5-fold) were observed in 30 top-ranked genes (Table 1) in microarray analysis including *Dnase2B, HspB1, γCry, MIP*, Beaded filament structural protein 1 (*Bfsp1*), and filensin that have been reported to be associated with cataract and lens development [21-25]. Furthermore, the expressions of six genes were upregulated (2.1 to 2.5-fold) in rat LECs from Cat+ compared to those from Cat-. Of the six genes, five genes were unidentified, and the other was Schlafen 4. However, the function of Schlafen4 in lens is not clear.

**Table 1:**
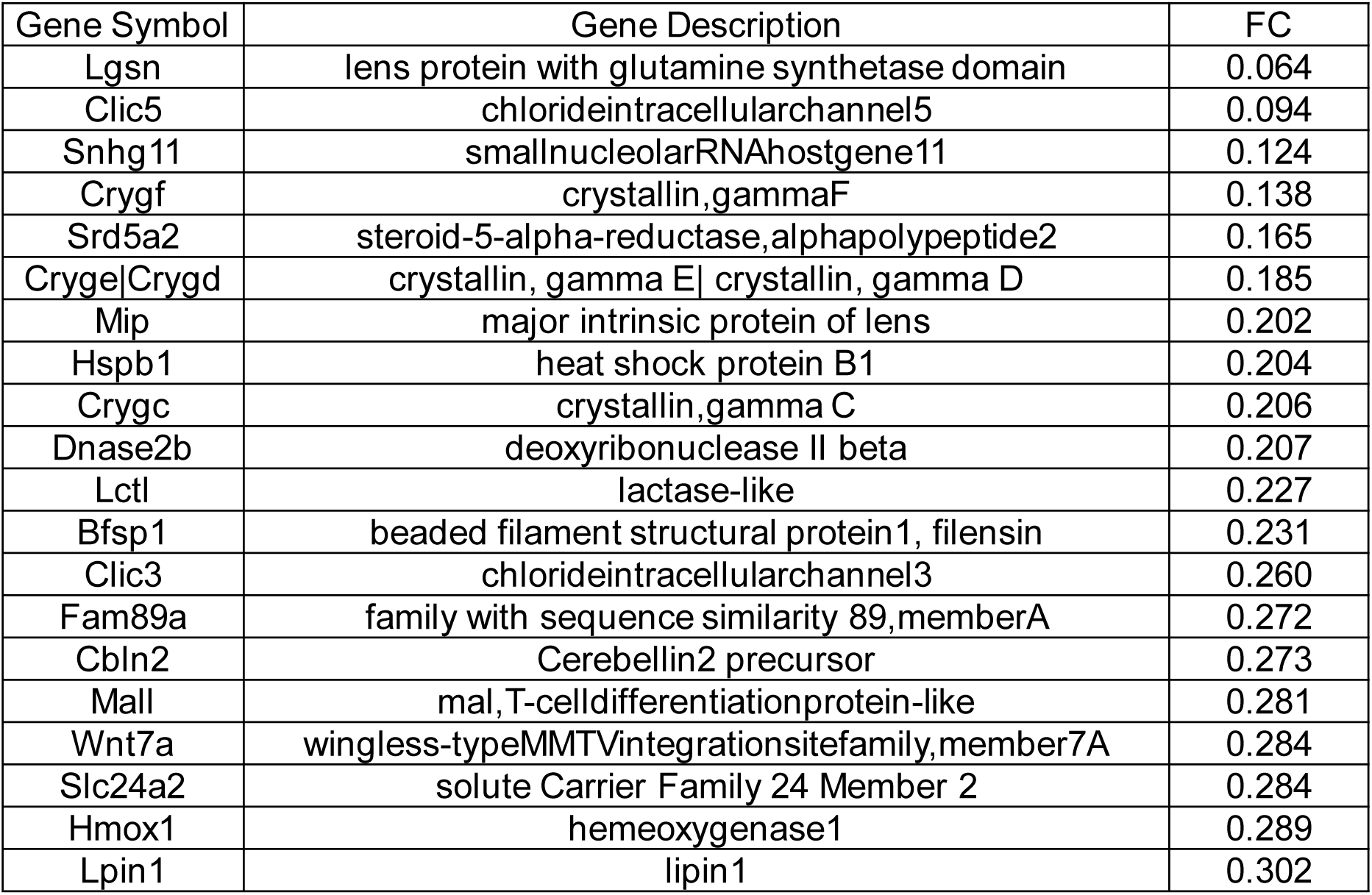
Lists of top 20 genes which are <0.5 down-regulated in LECs in 5-week-old Cat+ SCR compared to Cat-SCR.

**Fig. 2:**
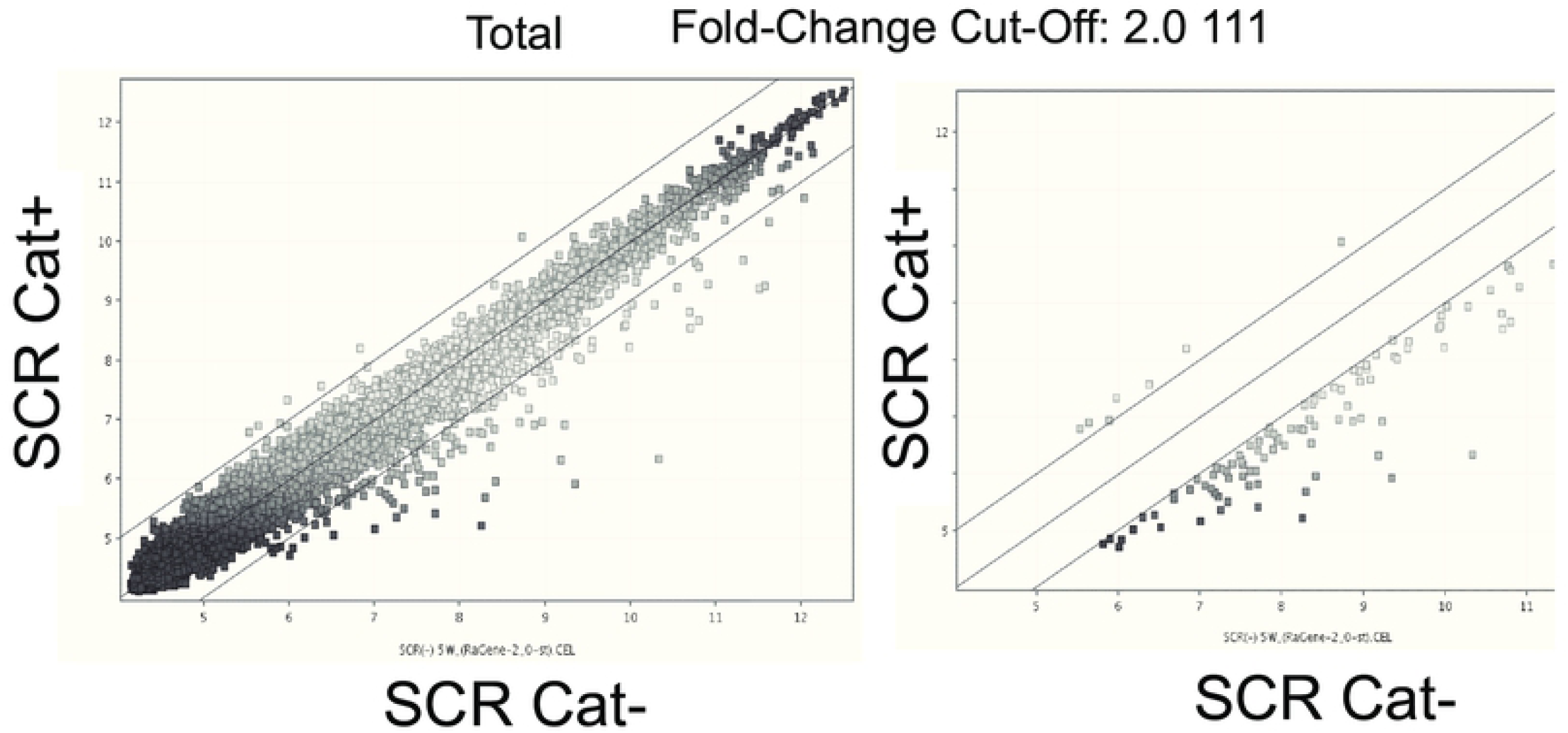
Analysis of gene expression profile. The scatter plots show a representative experiment using lens cDNAs from the Cat- and Cat+ SCRs at 5 weeks. Horizontal and vertical axes represent normal and experimentally generated signals on a logarithmic scale. The X-axis represents the log2 fold change and the dark vertical lines represent cut-offs at 2-fold decrease and increase.

### Validation of gene expression data using RT-qPCR

The data from the microarray experiment shown above revealed the downregulation of various genes during the progression of cataract. However, we observed that genes that belonged to the same class or family had similar expression patterns, indicating the possibility of cross-hybridization in microarray analysis. Therefore, we selected the following four genes: *Dnase2B, HspB1, γCry*, and *MIP* that showed significant changes in expression according to the microarray data, and accordingly, we validated these results using RT-qPCR Data from the RT-qPCR showed that *Dnase2B, Hspb1*, and *MIP* mRNA showed a significant downregulation in 5- and 10-week old Cat+ SCRs compared to Cat-SCRs (Fig. 3a, b, and c). However, the expression of *CryγD* mRNA was significantly downregulated only in 5-week-old Cat+ SCRs (Fig. 3d).

**Fig. 3:**
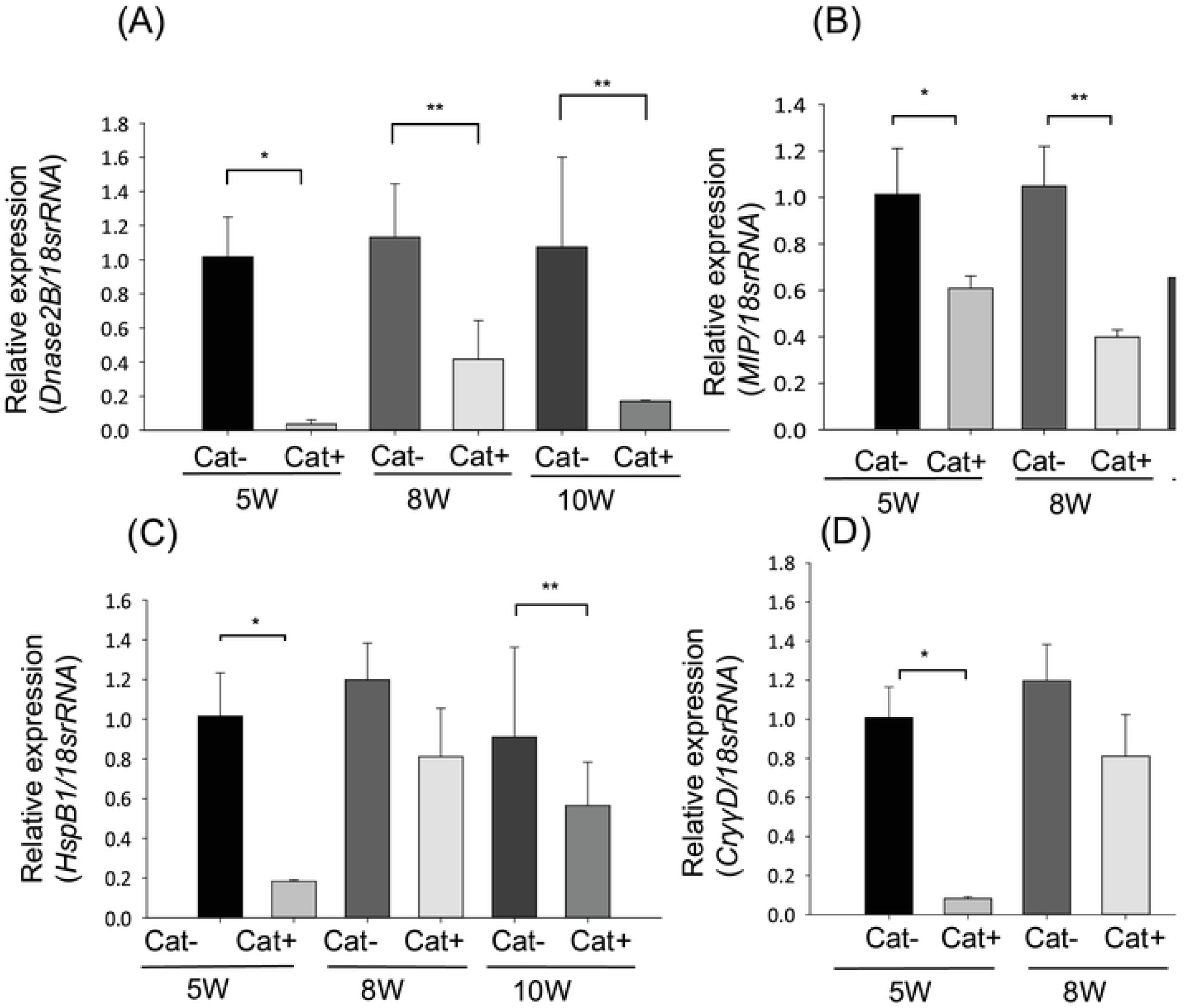
Expressions of *Dnase2B, MIP, HspB1* and *CryγD* mRNA in LECs from Cat-SCR and Cat+ SCR at 5, 8, and 10 weeks of age. A: Expression of *Dnase2B* mRNA; *p<0.02, **p<0.04. B: Expression of *MIP* mRNA.; *p<0.015, **p<0.001, ***p<0.04. C: Expression of *HspB1* mRNA.; *p<0.02, **p<0.04. D: Expression of CryγD mRNA.; *p<0.015, **p<0.001, ***p<0.04. Data are expressed as the mean ± standard deviation.

## Discussion

In this study, we comprehensively analyzed gene expression changes in the lens depending on the presence or absence of LSS deficiency in SCRs of different ages. SCRs with LSS deficiency gradually develop cataract and show mature cataract at 11 weeks of age. Lens opacity is not observed in 5-week-old SCRs with *Lss* mutations. In microarray analysis, 110 genes were downregulated with 0.5-fold change in Cat+ SCR at 5 weeks of age. It is speculated that carrying *Lss* mutation alters many gene expressions in the lens and induces lens opacity. In lens epithelial cells of SCRs with *Lss* mutations, the expression of many genes reported to be associated with cataract development was downregulated. In this study, we analyzed *Dnase2B, HspB1, γCry*, and *MIP* genes. We confirmed that expressions of *Dnase2B, HspB1*, and *MIP* mRNAs were significantly downregulated in the LECs of SCRs with *Lss* mutation before and after cataract onset. Furthermore, significant downregulation in the expression of *γCry* mRNA was observed in the LECs of SCRs with *Lss* mutation only before the onset of the disease.

In mice, DNase II-like-acid DNase (*DLAD*; *Dnase2B*) has been identified as a DNA-degrading enzyme that functions during lens enucleation [25]. Since the optimum pH for DLAD is acidic, it was suggested that the nucleus could be engulfed by lysosomes through autophagy and denucleated by the action of DLAD [25]. The lens consists of lens epithelial cells and differentiated lens fiber cells. In the process of fiber differentiation, intracellular structures such as the nucleus, mitochondria, and endoplasmic reticulum disappear and the lens becomes transparent. In the process of enucleation, the DNA that encodes the genetic information is degraded. For this enucleation process, DLAD plays an important role. In *DLAD* knock-out mice, the eye lens seemed to have developed normally; however, undegraded DNA was observed in the lens fiber that contributed to lens opacity. Thus, DLAD is necessary to maintain lens transparency and normal fiber differentiation. *Lss* mutation downregulates the expression of *DLAD* (*Dnase2B*), which could be attributed to cataract development in SCRs.

Heat shock protein beta-1 (HspB1) also known as heat shock protein 27 (Hsp27) is a protein that is encoded by the *HspB1* gene in humans. HspB1 is an ATP-independent molecular chaperone with a conserved αB-crystalline domain in the C-termini region [26]. Heat shock proteins (Hsp) play a central role in maintaining cellular homeostasis and altering protein folding, thereby protecting αA- and other crystalline proteins [27]. Further investigation of Hsp27 revealed that the protein responds to cellular oxidative and chemical stress conditions other than heat shock [27]. In the presence of oxidative stress, Hsp27 plays a role as an antioxidant, decreasing the ROS by raising levels of intracellular GSH [28, 29]. It has been reported that mutation in *HspB1* and/or αB-crystallin are responsible for the development of cataract [30] and are considered as major targets for the development of anti-cataract drugs [31]. We have previously reported that expression of Prdx6, an anti-oxidant protein, is decreased resulting in an increase in ROS in the LECs of SCRs [9]. Thus, *Lss* mutation may be involved in the decreased expression of anti-oxidant genes, causing oxidative stress and inducing cataract in SCRs.

MIP is a lens fiber major intrinsic protein, also known as Aquaporin 0, which is a water channel in lens fiber cells, facilitating the movement of water, gap junction channels, and solute transporters [32]. MIP plays a crucial role in regulating the osmolarity and homeostasis of the lens and stabilizing cell junctions in the lens nucleus. Currently, 12 mutations in *MIP* have been linked to autosomal-dominant cataracts in humans [22]. The decreased expression of *MIP* in SCR LECs may disrupt cellular water homeostasis and induce lens fiber swelling and vacuole formation observed in SCR lenses.

The nuclear region of the eye lens is particularly rich in the *γCry* protein, which is necessary to maintain structural and functional properties in the lens. *γCry* mRNA was downregulated in the LECs of Cat+ SCRs before the onset of lens opacity compared to Cat-rats. However, there was no significant change in expression of this gene after the onset of cataract. It is not clear why *γCry* mRNA expression did not decrease in cataractous lenses in SCRs. Further studies are required to understand this phenomenon.

Sterol, such as cholesterol in animals, is a compound that is important as a biosynthetic raw material for steroid hormones, vitamin D, and bile acids, in addition to its role in regulating membrane fluidity as a component of eukaryotic biological membranes [33]. Cholesterol synthesis is carried out through a process of approximately 30 enzymatic reactions using acetyl-CoA as a starting substrate. Lanosterol is the first sterol in the cholesterol synthesis pathway and LSS is an essential rate-limiting enzyme that functions as a downstream element in the lanosterol biosynthetic pathway [34], catalyzing the cyclization of the linear 2,3-monoepoxysqualene to cyclic lanosterol [34]. Previous studies have reported that LSS might play a significant role in oxidative stress and maintenance of lens transparency [35]. Congenital cataract with homozygous and heterozygous *Lss* mutations has been reported to affect the catalyzing functions of LSS [18, 19]. However, the relationship between LSS pathway and age-related cataract is unclear. Epidemiological studies have shown that individuals receiving statins, which are cholesterol synthesis inhibitors, have an increased risk of being diagnosed with cataracts [36, 37]. Recently, it has also been reported that lanosterol plays a preventive role in cataract formation, inhibiting lens opacity and reversing crystalline aggregation [18]. Additionally, intravitreal injection of lanosterol nanoparticles has been reported to rescue early stage of lens damage in SCRs [7]. Thus, synthesis of cholesterol by LSS is important to maintain lens transparency.

In conclusion, our study demonstrated that *Lss* mutations in SCRs downregulated the expressions of several genes associated with maintaining lens transparency and identified their relationship with cholesterol deficiency. Cholesterol and LSS in lens may be important to maintain normal lens homeostasis such as lens fiber differentiation, oxidative and heat stresses, and regulation of lens osmolarity to maintain lens transparency.

## Acknowledgments

This work was supported by grants from Japan Society for the Promotion of Science (JSPS) KAKENHI Grant Numbers JP 17K11470 (to EK) and National EYE Institute, National Institute of Health (NIH) (EY024589) to (DPS). We are thankful to the National Bio Resource Project-Rat (http://www.anim.med.kyoto-u.ac.jp/NBR/) for providing the rat strains (SCR/Sscr: NBRP Rat No. 0823). We would like to thank Editage (www.editage.com) for English language editing.

